# Integrative analyses of the RNA modification machinery reveal tissue- and cancer-specific signatures

**DOI:** 10.1101/830968

**Authors:** Oguzhan Begik, Morghan C. Lucas, Huanle Liu, Jose Miguel Ramirez, John S. Mattick, Eva Maria Novoa

## Abstract

**Background:** RNA modifications play central roles in cellular fate and differentiation. These features have placed the epitranscriptome in the forefront of developmental biology and cancer research. However, the machinery responsible for placing, removing and recognizing more than 170 RNA modifications remains largely uncharacterized and poorly annotated, and we currently lack integrative studies that identify which RNA modification–related proteins (RMPs) may be dysregulated in each cancer type.

**Results:** Here we have performed a comprehensive annotation and evolutionary analysis of human RMPs as well as an integrative analysis of their expression patterns across 32 tissues, 10 species and 13,358 paired tumor-normal human samples. Our analysis reveals an unanticipated heterogeneity of RMP expression patterns across mammalian tissues, with a vast proportion of duplicated enzymes displaying testis-specific expression, suggesting a key role for RNA modifications in sperm formation and possibly intergenerational inheritance. Moreover, through the analysis of paired tumor-normal human samples we uncover many RMPs that are dysregulated in various types of cancer, and whose expression levels are predictive of cancer progression. Surprisingly, we find that several commonly studied RNA modification enzymes such as METTL3 or FTO, are not significantly up-regulated in most cancer types, once the sample is properly scaled and normalized to the full dataset, whereas several less-characterized RMPs, such as LAGE3 and HENMT1, are dysregulated in many cancers.

**Conclusions:** Our analyses reveal an unanticipated heterogeneity in the expression patterns of RMPs across mammalian tissues, and uncover a large proportion of dysregulated RMPs in multiple cancer types. We provide novel targets for future cancer research studies targeting the human epitranscriptome, as well as foundations to understand cell type-specific behaviours that are orchestrated by RNA modifications.

## BACKGROUND

Technological advancements have revolutionized our understanding of RNA modifications, which can occur by post-transcriptional removal (by deamination, often called ‘RNA editing’) or by the addition of chemical side groups on the ribose or base moieties. These chemical entities, collectively known as the ‘epitranscriptome’ [1], not only occur in tRNAs and rRNAs, where they were first identified and have traditionally been studied, but also in other molecules, such as mRNAs, long noncoding RNAs, piRNAs and miRNAs [2–6]. A number of studies have shown that RNA modifications can profoundly affect central biological processes, including cell fate [7], sex determination [8,9], maternal-to-zygotic transition [10] and the circadian clock [11]as well as plant developmental timing, morphogenesis and flowering [12]. Furthermore, dysregulation of their activity has been associated with more than 100 different human diseases [13–17]. At a molecular level, modifications can affect the fate and function of the RNA molecules that contain them, including turnover rates [18–20], translation efficiency [21,22] and subcellular localization [23], amongst others.

Over 170 different RNA modifications are known to decorate RNA molecules [24]. In the last few years, a vast amount of efforts have been devoted to functionally dissecting the biological role of N6-methyladenosine (m^6^A), the most prevalent internal RNA modification found in human mRNAs. M^6^A is placed by a multicomponent transferase complex, in which methyltransferase 3 (METTL3) acts as the catalytic subunit [25,26]. Moreover, m^6^A modifications can be reversed by the activity of m^6^A demethylases or ‘erasers’, namely the fat mass and obesity-associated protein (FTO) [27] and the alkB homolog 5 (ALKBH5), although recent studies have suggested that only the latter can demethylate m^6^A marks [28]. Mechanistically, m^6^A modifications can alter mRNA splicing [29–31], cause mRNA decay [20] and affect translation [2,32,33]; thus, they can govern major cellular processes including cellular fate [34,35], stress responses [2] and differentiation programs [36]. These features have set out m^6^A marks, and more specifically, their ‘readers’, ‘writers’ and ‘erasers’, as promising drug targets for multiple diseases, including cancer [37–40]. However, the functional characterization of the majority of RNA modifications still remains an uncharted territory.

Insights into the physiological roles of specific RNA modification-related proteins (RMPs) have mostly come from naturally occurring phenotypes or diseases associated with their loss of function [13–17]. However, a systematic annotation and characterization of RNA modification RMPs across human tissues, cell types, and disease states is currently lacking.

Here we have compiled and analyzed the evolutionary history of 90 RNA modification writers as well as the gene expression patterns of 146 human RMPs (**Table S1**) from 32 tissues, 10 species and 13,358 tumor-normal samples. Our analyses revealed that many RMPs display restricted gene expression patterns and/or are dysregulated in specific types of cancer. Specifically, we found that a vast proportion of RNA modification ‘writers’ have undergone duplications (84%), and that these were typically accompanied by a change in their RNA target specificity and/or tissue expression patterns (82%). We observed that the most frequent change in tissue specificity is the acquisition of restricted testis-specific expression, suggesting that a significant portion of the human RNA modification machinery is likely devoted to sperm formation and maturation. We also found that 27% of human RMPs are significantly dysregulated in cancers, and identified several dysregulated RMPs whose expression is strongly correlated with cancer prognosis. Overall, our work reveals an unanticipated heterogeneity of RMP expression across both normal and malignant cell types, and points towards several less-characterized RMPs, such as HENMT1 or LAGE3, as promising drug targets for antitumor therapies.

## RESULTS

### Comprehensive annotation and evolutionary analysis of RNA modification writers

To reveal the evolutionary history of the RNA modification machinery, we first compiled and manually curated a list of human RMPs (**Table S1**, see also *Methods*). Due to the wide chemical variety of RNA modifications, we restricted our evolutionary analysis to the catalytic domain of three major RNA modification ‘writer’ (RMW) classes: i) methyltransferases, ii) pseudouridylases and iii) deaminases. For each annotated RMW [13,41,42], Pfam domains of the catalytic domain were extracted and used as input for HMM-based searches against the human proteome. This resulted in a total of 90 human RMWs, doubling the amount annotated human RMWs in other resources [41]. To determine the evolutionary history and identify duplication events occurred in each family, ortholog proteins from representative species were retrieved (see *Methods*), and phylogenetic trees were built to identify the number of duplications occurred within each family. Overall, our analysis identified 46 duplication events (**Figure 1A**), which have mainly occurred in the base of Eukaryota, Metazoa and Vertebrata (**Figure 1B,** see also **Table S2).**

**Figure 1.**
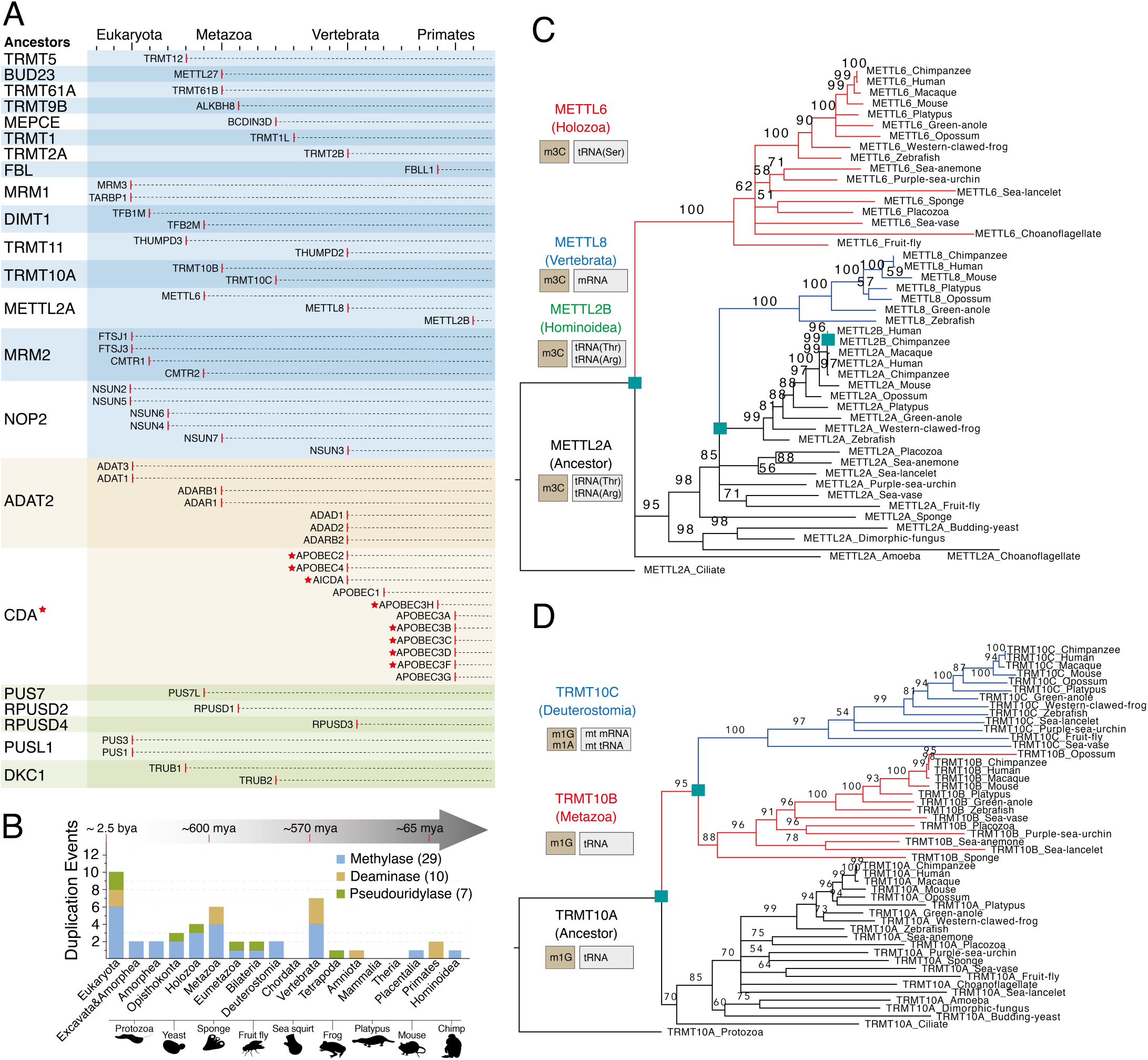
Evolutionary analysis of RNA modification ‘writers’. **(A)** Detailed overview of the evolutionary history of RMW duplications during eukaryotic evolution. Red stars indicate that proteins do not target RNAs but they are in the same family with an RNA writer protein. Red lines indicate the time where the enzyme has appeared. **(B)** Histogram of RMW duplication events throughout eukaryotic evolution. Duplication events were inferred by using multiple sequence alignments, coupled to maximum likelihood tree generation, for each family. **(C and D)** Maximum Likelihood phylogenetic trees of the methyltransferase family METTL2A/2B/6/8 (C) and TRMT10A/B/C (D). Cyan squares indicate the node where the duplication occurred. Numbers shown on the branches indicate bootstrapping values.

We find that duplications are often accompanied by changes in substrate specificity (**Figures 1C,D**), at least in those RMWs where the substrate specificity has been reported. One such case is the family of 3-methylcytosine (m^3^C) RNA methyltransferases, where the ancestral protein methyltransferase-like protein 2 (METTL2) modifies both tRNA^Arg^ and tRNA^Thr^, whereas its paralog enzymes, METTL6 and METTL8, methylate tRNA^Ser^ and mRNA, respectively [43] (**Figure 1C**). Similarly, we find that the N1-methylguanosine (m^1^G) methyltransferases TRMT10A and TRMT10B modify tRNAs in position m^1^G9 [44], whereas its paralog TRMT10C has been reported to place N1-methyladenosine (m^1^A) in mitochondrial tRNAs and mRNAs [3], in addition to m^1^G in tRNAs (**Figure 1D**).

### Heterogeneity of expression patterns among duplicated RMPs is conserved across species

We then wondered whether duplicated RMPs might have acquired distinct tissue expression patterns - and possibly, different functions - than the ancestral gene. To test this, we examined the heterogeneity of RMP expression patterns across tissues in human and mice, using publicly available RNASeq datasets [45–47] (**Figure 2A,** see also **Figures S1A** for gene-labelled heatmaps). For each gene and tissue, we computed ‘tissue specificity (TS) scores’ [48], which is defined as the deviation of gene expression levels in a given tissue, relative to the average expression across all tissues (see *Methods*). Using this approach, we found that testis is the most distinctive tissue in terms of RMP gene expression patterns, both in human and mouse (**Figure 2A,B)**. This was due to several RMPs being quasi-exclusively expressed in testis (e.g. ADAD1, ADAD2), but also to several RMPs whose expression levels are significantly increased this tissue (e.g. FBLL1, HENMT1, NSUN7). In contrast, other tissues such as colon did not display none or few tissue-enriched RMPs (**Figure 2B,** see also **Figure S1B**). Moreover, we found that RMP tissue expression patterns were largely conserved in both mouse and human (**Figure 2C,** see also **Table S3**),

**Figure 2.**
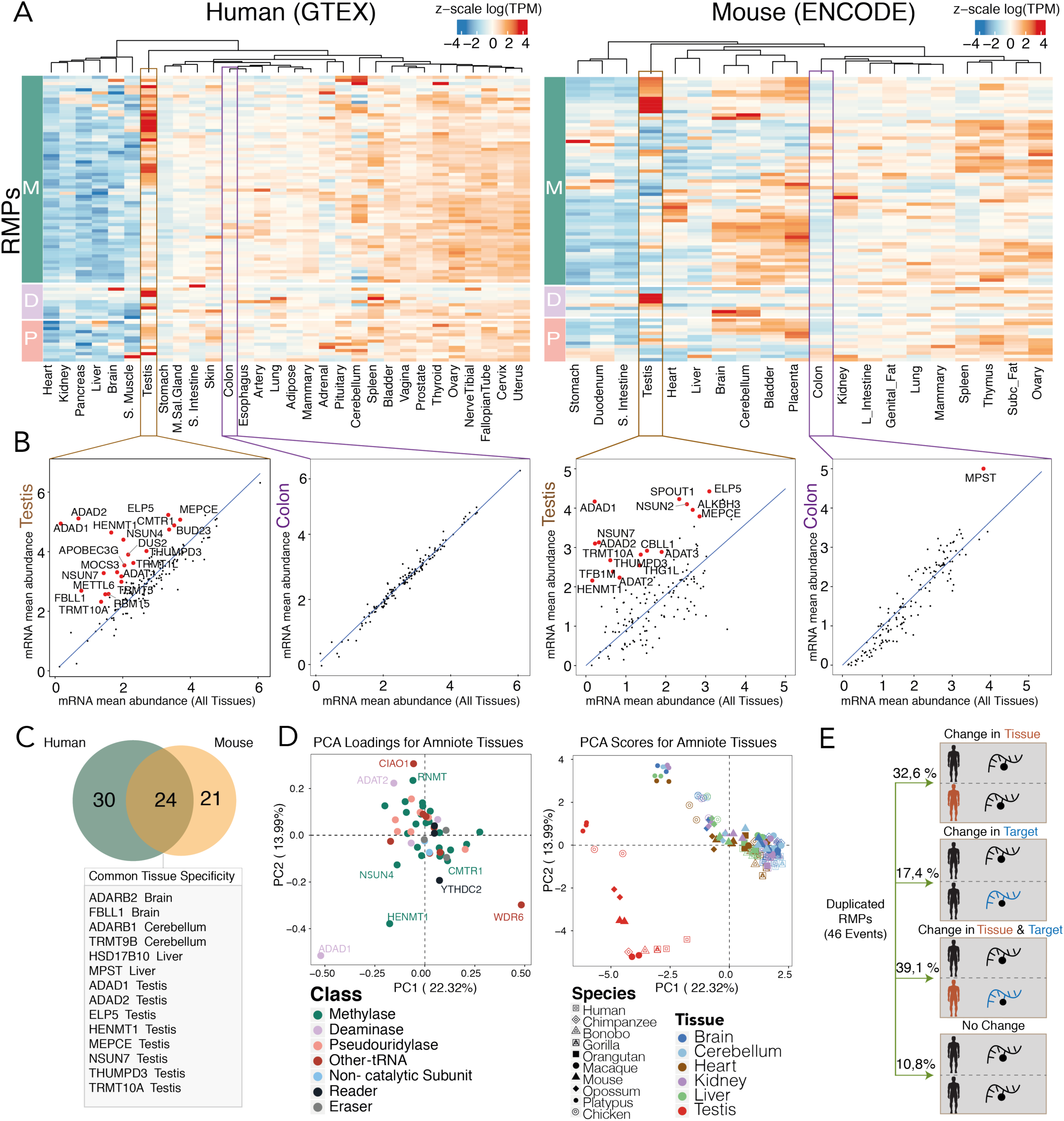
Analysis of RMP tissue specificity expression in different species. **(A)** Heatmap of z-scaled log(TPM) values of catalytic RNA writer proteins (M: methyltransferases; D: deaminases; P: pseudouridylases) throughout human and mouse tissues. In both human and mouse, testis has the most distinct RMP expression pattern in which many genes show really high expression, whereas other tissues such as colon shows moderate expression level of RMPs. **(B)** Scatter plots depicting tissue-specificity analysis, which have been computed by representing the RMP mRNA expression values in a given tissue (y axis) relative to the mean mRNA abundance in all tissues (x axis). Scatter plots show that testis has a significant number of tissue-specific genes in both human and mouse, while colon shows no tissue-specific genes in human and only one in mouse. Tissue-specific genes are labelled in red. **(C)** Venn diagram of the conservation of tissue specificity between human and mouse. Out of 24 common tissue-specific genes, 14 of them are specifically expressed in the same tissue. **(D)** Principal Component Analysis of amniote tissues based on the log(RPKM) mRNA expression of their RMPs. The loadings plot (left) shows the contribution of each RMP to the clustering of amniote tissues. The scores plot (right) shows the clustering of each tissue, where testis tissue (in red) is the main contributor to the variance of the data, and is found apart from the rest of the amniote tissues for every given species. **(E)** Schematic representation of the fate of the 46 RMW duplication events shown in Figure 1, showing that 89% of them suffered a change in their tissue and/or target specificity.

We then extended our analysis to additional amniote species, finding that testis was also the main outlier in terms of RMP expression patterns in all species analyzed, supporting the notion that testis-specific RMP functionalities are conserved (**Figure 2D**, see also **Figure S2**). Overall, we found that 89% of RMP duplication events were often followed by a change in tissue specificity (32.6%), target specificity (17.4%) or both (39.1%) (**Figure 2E**, see also **Table S4**), with a major over-representation of acquisition of testis-specific gene expression.

### Testis-specific RMPs are mainly expressed during meiotic stages of spermatogenesis

The process of sperm formation, termed spermatogenesis (**Figure 3A**), is a highly-specialized differentiation process in which transcriptional, post-transcriptional and translational regulation are highly orchestrated [49–52]. RNA modifications can influence pre-mRNA splicing, mRNA export, turnover, and translation, which are controlled in the male germline to ensure coordinated gene expression [35]. Recent works have shown that m6A depletion in mice dysregulates translation of transcripts that are required for spermatogonial proliferation and differentiation [34]. Moreover, m^5^C modifications have been shown to be essential for transmission of diet-induced epigenetic information across generations in the epididymis [53]. However, whether additional RNA modifications may be involved in such orchestration is largely unknown.

**Figure 3.**
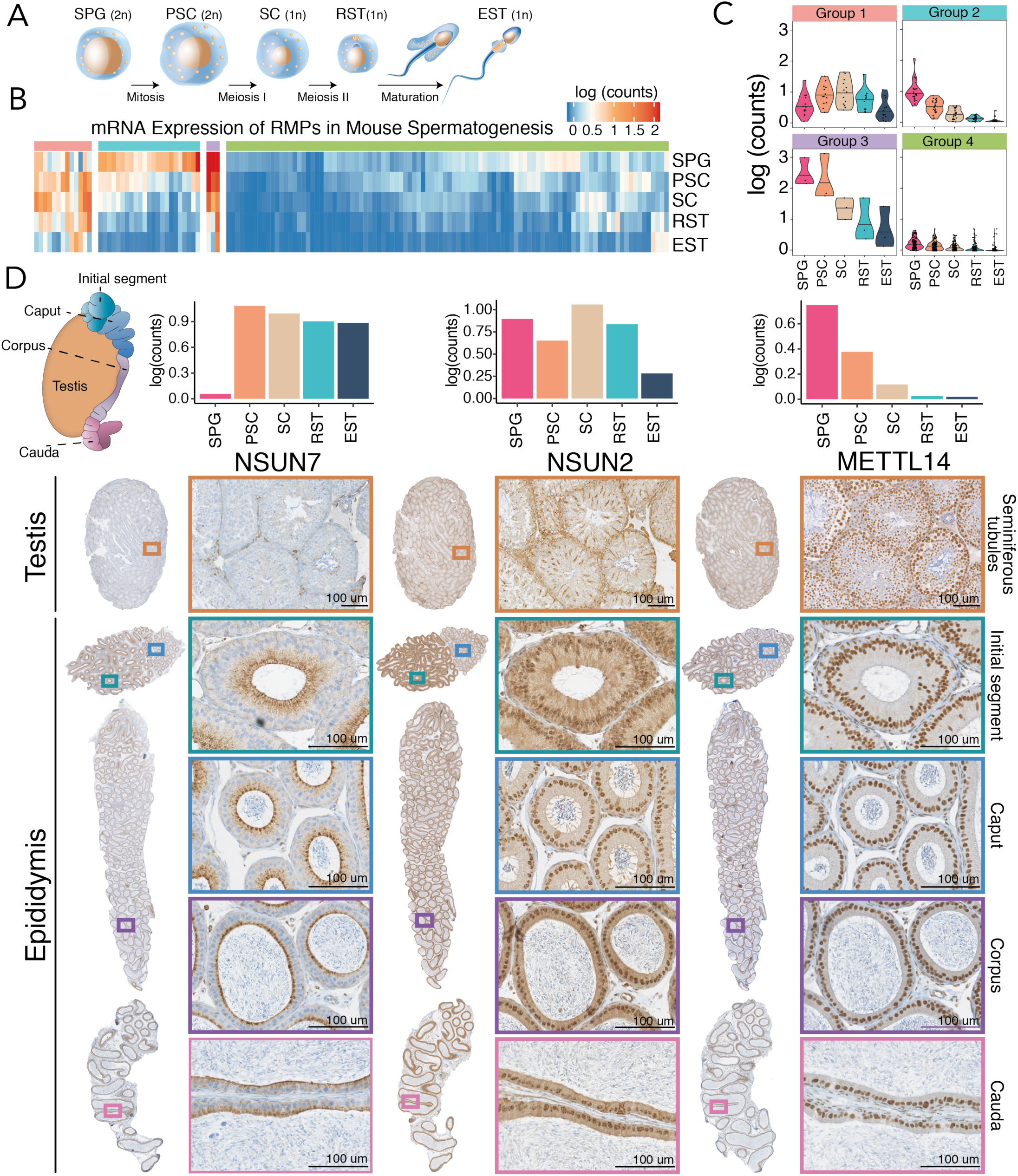
Analysis of RMP gene expression during spermatogenesis. **(A)** Schematic representation of the four main phases of spermatogenesis: i) mitotic division of spermatogonia (SPG) into primary spermatocytes (PSC) ii) meiotic division of PSCs into secondary spermatocytes (SC), iii) meiotic division SCs into round spermatids (RST) and iv) spermiogenesis, in which round spermatids (RST) mature into elongated spermatids (EST). **(B)** Heatmap of RMP expression levels of in mouse testis. RMPs were clustered into 4 groups based on k-means analysis of their normalized average mRNA expression values. **(C)** Violin plots of the expression patterns of each of the 4 identified clusters **(D)** RNA median expression barplot and immunohistochemistry of NSUN7, NSUN2 and METTL14, depicting distinct protein expression levels along the different sections of the testis and epididymis, as well as different subcellular localisations. Brown color indicates a specific staining of the antibody whereas blue represents hematoxylin counterstain.

To identify at which stage of sperm formation and maturation testis-specific RMPs are involved in, we gathered publicly available single-cell RNA sequencing data from mouse testis [54] (**Figure 3B**, see also **Figure S3** for gene-labelled heatmap). We first classified RMPs based on their gene expression patterns (see *Methods*), identifying four main expression patterns: (i) high expression only during meiotic stages (spermatocytes and spermatids); (ii) high expression only during mitotic stages (spermatogonia); (iii) high expression in both mitotic and meiotic stages (spermatogonia, spermatocytes, spermatids), although decreased in the latter; and (iv) low expression throughout spermatogenesis (**Figure 3B-C**). We find that the majority of RMPs, including those involved in placing, removing m6A (VIRMA, YTHDC2, YTHDF2, ALKBH5, METTL14, METTL3) are highly expressed in spermatogonial cells, whereas their expression rapidly drops as the spermatogenic process begins (**Figure 3B-C**, see also **Figure S3C-D**).

Interestingly, this is not the case for all RMPs, such as m^5^C methyltransferase NSUN7, which is not expressed in early stages of spermatogenesis, but whose expression levels are drastically increased in spermatocytes and spermatids (**Figure S3A, C**). Similarly, the testis-specific adenosine deaminase ADAD1 is not expressed in early stages of spermatogenesis, but its expression levels are greatly increased in meiotic stages. Depletion of NSUN7 or ADAD1 are known to cause infertility [55,56], suggesting that RMPs that are selectively expressed in meiotic stages of spermatogenesis are essential for proper sperm formation and/or maturation. However, the molecular mechanisms behind these infertility phenotypes are largely uncharacterized.

We then investigated whether specific RMPs also showed increased expression patterns in epididymis, relative to other tissues (**Figure S3D**). Our analysis identified two RMPs as epididymis-enriched: (i) TRDMT1-also known as DNMT2-, a N5-methyladenosine (m^5^C) methyltransferase modifying position 38 in specific tRNAs [57], and (ii) METTL1, a N7-methylguanosine (m^7^G) tRNA methyltransferase, which has been recently shown to act not only on tRNAs, but also on miRNAs, promoting their maturation [58]. Interestingly, TRDMT1 has been shown to play a major role in the transmission of paternal epigenetic information across generations [53]; however, whether METTL1 is involved in the transmission of such information is yet to be determined.

### Immunohistochemistry reveals heterogeneity in RMP expression patterns along the epididymis

It is well established that mRNA levels do not always correlate well with protein levels [59]. Thus, to assess whether our findings would hold at the protein level, we performed immunohistochemistry in both testis and epididymis to characterize the expression patterns of 4 RMPs at the protein level: (i) NSUN7, a putative m^5^C methyltransferase that has been shown to affect sperm motility [56,60]; (ii) NSUN2, an m^5^C tRNA methyltransferase involved in sperm differentiation [61]; (iii) METTL14, a component of the m^6^A methyltransferase complex, which has been shown to be dynamically regulated during spermatogenesis [34], and (iv) HENMT1, a piRNA 2’-O-methyltransferase responsible for transposon silencing during spermatogenesis [5] (**Figure 3D**, see also **Figure S3E**).

We found that NSUN7 is most highly expressed in chromatoid bodies of spermatocytes, as well as in the initial segment and caput regions of the epididymis, in agreement with its role in the acquisition of sperm motility [56,60,62,63] (**Figure 3D,** left panels). Intriguingly, NSUN7 displayed vesicular subcellular localisation in the epithelial cells of epididymal ducts, with significant accumulation in the apical surface. It is yet to be determined how NSUN7 depletion causes defects in sperm motility, as well as which are the targets of NSUN7 in testis and epididymis tissues. On the other hand, NSUN2 displayed high expression levels in the nucleolus of the spermatogonial cells and spermatocytes (**Figure 3D,** middle panels), supporting its role in sperm differentiation [61]. We also observed that NSUN2 is highly expressed in the initial segment of the epididymis, with decreased expression in the remaining epididymal sections.

METTL14 was also found highly expressed in early spermatogenesis and down-regulated during the subsequent stages (**Figure 3D,** right panels), in agreement with the dynamic regulation of m^6^A levels during spermatogenesis [34]. Finally, HENMT1 is mainly expressed in round and elongating spermatocytes, where it is accumulated in the chromatoid bodies (**Figure S3E**). Chromatoid bodies are known to contain large amounts of piRNAs, in agreement with the role that HENMT1 plays in piRNA stability during spermatogenesis [5]. Overall, our analyses showed that RMPs are dynamically expressed during spermatogenesis and during sperm maturation, and that, for the four genes investigated, protein expression patterns were in agreement with mRNA expression.

### Analysis of RMP expression in tumor-normal paired human samples reveals heterogeneity in RMP dysregulation across cancer types

Due to their ability to modulate RNA metabolism and influence protein synthesis rates, RNA modifications have recently emerged as important regulators of cancer [64–66]. Several studies have shown that modulation of the RNA modification machinery can decrease the expression of specific oncogenes [36,67]. For example, in the case of glioblastoma, treatment with an FTO inhibitor was shown to decrease the expression levels of certain oncogenes [65]. Similarly, tRNA modifying enzymes NSUN2 and METTL1 can affect chemotherapy sensitivity by changing the methylation states of certain tRNAs [68]. Thus, understanding which epitranscriptomic players are dysregulated in each tumour type is essential to guide the research for future anticancer therapies targeting this regulatory layer.

To this end, we performed an integrative analysis of RMPs gene expression across 13,358 tumor-normal paired human samples gathered from publicly available datasets [69], which included 28 different cancer types (**Table S5**). Firstly, we compared the expression patterns between paired tumor-normal samples by measuring the log2 fold changes of median gene expression between tumor and normal paired samples, for each RMP and cancer type (**Figure S4** and *Methods*). We found that certain cancer types, such as pancreatic adenocarcinoma (PAAD) and acute myeloid leukemia (LAML) showed significant dysregulation of a vast proportion of RMPs (**Figure S4**). Surprised by this result, we wondered whether these global up/down-regulation patterns could in fact be an artefact generated by the use of external datasets. Indeed, certain TCGA cancer types do not have real ‘matched’ tumor-normal data readily available, and often employ data from other publicly available datasets (e.g. GTEx) as ‘normal’ human tissue (**Table S5**).

To address this issue, we extracted the gene expression levels of all genes - not just RMPs - for each cancer type, finding that certain cancer types that employ GTEx data as source of ‘normal’ human tissues, such as LAML, display low Pearson correlation values between matched tumor-normal samples (r^2^= 0.86), compared to those observed in other cancer types such as prostate adenocarcinoma (PRAD) (r^2^=0.98) **(Figure S6**). Thus, to identify which RMPs were significantly dysregulated in each cancer type, we computed ‘dysregulation scores’ [48], which take into account the global variance of the tumor-normal paired data, for each cancer type (**Figure 4A**). We considered an RMP as dysregulated in a given cancer type if its dysregulation score was higher than 2.5 standard deviations (SD) relative to the linear fit to the gene expression in the matched normal tissue (see *Methods*). Using this strategy, we identified a total of 40 RMPs which are dysregulated in at least one cancer type (**Table 1,** see also **Table S6**). Moreover, we find that the ‘global’ up/down-regulation patterns found using log2 fold change comparisons are not further observed, suggesting that these results were in fact artefacts caused by the lack of proper ‘matched’ normal tissues for certain cancer types.

**Table 1.** List of significantly dysregulated RMPs identified using dysregulation score-based analysis

| Short Name | Cancer type | Up-regulated RMPs | Down-regulated RMPs |
| --- | --- | --- | --- |
| ACC | Adrenocortical carcinoma | BUD23, FBL, LAGE3, TRMT112, VIRMA | ADAT2, APOBEC3G, TARBP1 |
| BLCA | Bladder Urothelial Carcinoma | HENMT1 | ADARB1 |
| BRCA | Breast invasive carcinoma | LAGE3 | TRMT9B |
| CESC | Cervical squamous cell carcinoma and endocervical adenocarcinoma | APOBEC3A, HENMT1 | - |
| COAD | Colon adenocarcinoma | APOBEC1, DKC1, HENMT1 | ADARB1 |
| ESCA | Esophageal carcinoma | HENMT1, RBM15, ZC3H13 | ADAD2, METTL17 |
| GBM | Glioblastoma multiforme | APOBEC3G, HSD17B10 | ADARB2, FBLL1, TARBP1 |
| HNSC | Head and Neck squamous cell carcinoma | HENMT1 | - |
| KICH | Kidney Chromophobe | ISCU | - |
| KIRC | Kidney renal clear cell carcinoma | APOBEC3G | - |
| KIRP | Kidney renal papillary cell carcinoma | METTL27 | - |
| LAML | Acute Myeloid Leukemia | APOBEC3A, HENMT1 | - |
| LGG | Brain Lower Grade Glioma | HSD17B10, TRMT5 | ADARB1, HENMT1, TARBP1 |
| LIHC | Liver hepatocellular carcinoma | LAGE3, METTL27, TARBP1 | ADAT2, METTL17, NSUN6, TRMT11, ZC3H13 |
| LUAD | Lung adenocarcinoma | LAGE3, METTL1, TFB2M | ADARB1, APOBEC3A, TRMT9B |
| LUSC | Lung squamous cell carcinoma | DKC1, FBL, LAGE3, METTL1, METTL8 | ADARB1, TRMT9B |
| OV | Ovarian serous cystadenocarcinoma | - | YTHDC2 |
| PAAD | Pancreatic adenocarcinoma | APOBEC1, APOBEC3G, HENMT1 | QTRT1, WDR6 |
| PCPG | Pheochromocytoma and Paraganglioma | FBLL1 | MPST |
| PRAD | Prostate adenocarcinoma | - | APOBEC3G, METTL17 |
| READ | Rectum adenocarcinoma | APOBEC1, DKC1, HENMT1 | ADARB1 |
| SARC | Sarcoma | HENMT1 | ADARB1, ISCU, SERGEF |
| SKCM | Skin Cutaneous Melanoma | APOBEC3G | - |
| STAD | Stomach adenocarcinoma | HENMT1 | METTL17, TRMT2A, WDR6 |
| TGCT | Testicular Germ Cell Tumors | EMG1, FBL, HNRNPC, HSD17B10, LAGE3, METTL9 | ADAD1, ADAD2 |
| THCA | Thyroid carcinoma | LAGE3, TRMT112 | ADAT2, METTL17, TRMT9B, TRMU |
| UCEC | Uterine Corpus Endometrial Carcinoma | LAGE3 | ADARB1 |
| UCS | Uterine Carcinosarcoma | FBL, HENMT1, HSD17B10, LAGE3, TRMT112 |  |

**Figure 4.**
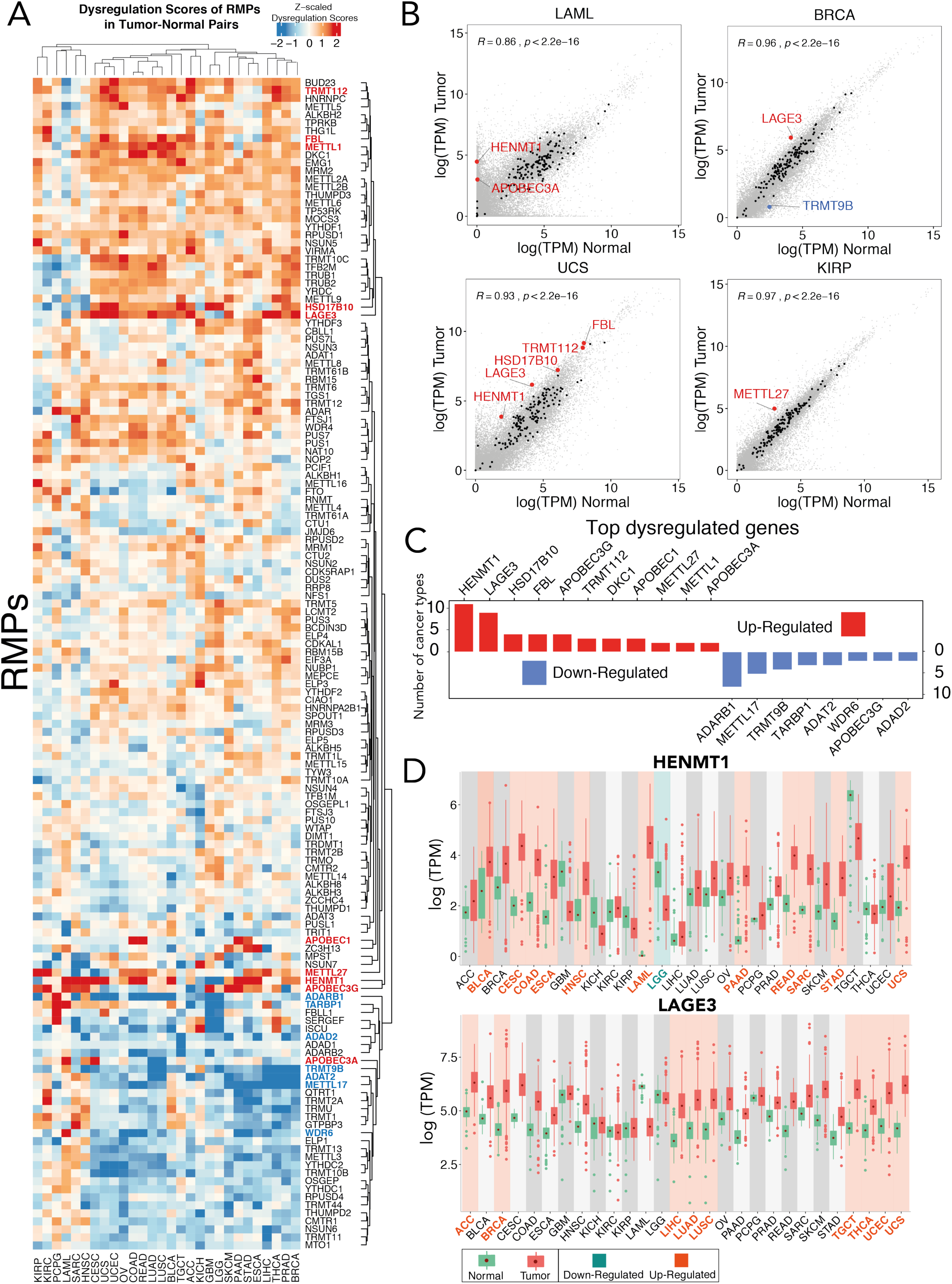
Expression analysis of RMPs in human tumor-normal paired samples. **(A)** Heatmap of z-scaled dysregulation scores of RMPs in tumor-normal paired samples, across 28 cancer types. Positive (red) values indicate up-regulation in tumor, whereas negative (blue) values indicate down-regulation. Genes labeled as red (up-regulated) and blue (down-regulated) represent top significantly dysregulated genes, which are also individually listed in panel C. **(B)** Scatter plot comparing RMP expression levels of matched tumor-normal samples, for the following cancer types: LAML (Acute Myeloid Leukemia) and UCS (Uterine Carcinosarcoma) BRCA (Breast invasive carcinoma) and KIRP (Kidney renal papillary cell carcinoma). Values represent median log(TPM) across all patients. Black data points indicate the expression of RMPs, where dysregulated genes are highlighted in red (up-regulated) or blue (down-regulated). Non-RMP genes are depicted in grey. **(C)** Barplot illustrating the number of cancer types in which significantly dysregulated genes highlighted in red (up-regulated) or blue (down-regulated). Only RMPs that are dysregulated in more than 2 cancer types are shown. For the full list of dysregulated RMPs, see Table 1. **(D)** Boxplots of log(TPM) mRNA expression values of HENMT1 (upper panel) and LAGE3 (bottom panel) across all 28 cancer types analyzed in this work. Green box plots represent normal samples, whereas red box plots represent tumor samples. Tumor-normal pairs highlighted in cyan represent cancer types in which the RMP is significantly down-regulated, whereas those highlighted in orange represent those cancer types in which the RMP is up-regulated. Error bars represent standard deviation of mRNA expression levels across patients. Each data point represents a different patient sample. Abbreviations: ACC (Adrenocortical carcinoma), BLCA (Bladder Urothelial Carcinoma), BRCA (Breast invasive carcinoma), CESC (Cervical squamous cell carcinoma and endocervical adenocarcinoma), COAD (Colon adenocarcinoma), ESCA (Esophageal carcinoma), GBM (Glioblastoma multiforme), HNSC (Head and Neck squamous cell carcinoma), KICH (Kidney Chromophobe), KIRC (Kidney renal clear cell carcinoma), KIRP (Kidney renal papillary cell carcinoma), LAML (Acute Myeloid Leukemia), LGG (Brain Lower Grade Glioma), LIHC (Liver hepatocellular carcinoma), LUAD (Lung adenocarcinoma), LUSC (Lung squamous cell carcinoma), OV (Ovarian serous cystadenocarcinoma), PAAD (Pancreatic adenocarcinoma), PCPG (Pheochromocytoma and Paraganglioma), PRAD (Prostate adenocarcinoma), READ (Rectum adenocarcinoma), SARC (Sarcoma), SKCM (Skin Cutaneous Melanoma), STAD (Stomach adenocarcinoma), TGCT (Testicular Germ Cell Tumors), THCA (Thyroid carcinoma), UCEC (Uterine Corpus Endometrial Carcinoma), UCS (Uterine Carcinosarcoma).

### Dysregulation score analyses of tumor-normal paired human samples identify LAGE3 and HENMT1 as top-ranked dysregulated RMPs

We then asked whether specific RMP genes were recurrently up- or down-regulated in multiple cancer types, as these could constitute promising drug targets that could be used to treat diverse cancer types. We identified 11 RMPs that were up-regulated in two or more cancer types, as well as 8 RMPs which were consistently down-regulated in at least 2 cancer types (**Figure 4C,** see also **Table 1**). We found that the most frequently up-regulated RMP was HENMT1 (**Figure 4D**), a piRNA 2’-O-methyltransferase which is highly expressed in gonadal cells, involved in transposable element (TE) mutagenesis protection [5,70,71]. Whether the global up-regulation of HENMT1 in cancer samples might be contributing to increased TE mutagenesis is currently unknown.

The second most frequently up-regulated RMP was the L antigen family member 3 (LAGE3), a component of complex responsible for formation of N6-threonylcarbamoyladenosine (t^6^A) in position 37 of tRNAs (**Figure 4D**). Interestingly, this modification is found in the anticodon stem-loop of many tRNAs decoding ANN codons [72], and has been shown to affect both translation accuracy as well as efficiency[73].

Finally, we asked whether the expression levels of RMPs might be correlated with cancer prognosis. We identified 283 cases where RMP expression patterns are significantly associated with patient survival (**Figure 5A,** see also **Table S7**). For example, we found that high NSUN5 expression levels in glioblastoma (GBM) are correlated with poor prognosis, in agreement with a recent study [74]. Similarly, our work revealed BUD23 expression to be correlated with cancer survival, in agreement with another recent study [75].

**Figure 5.**
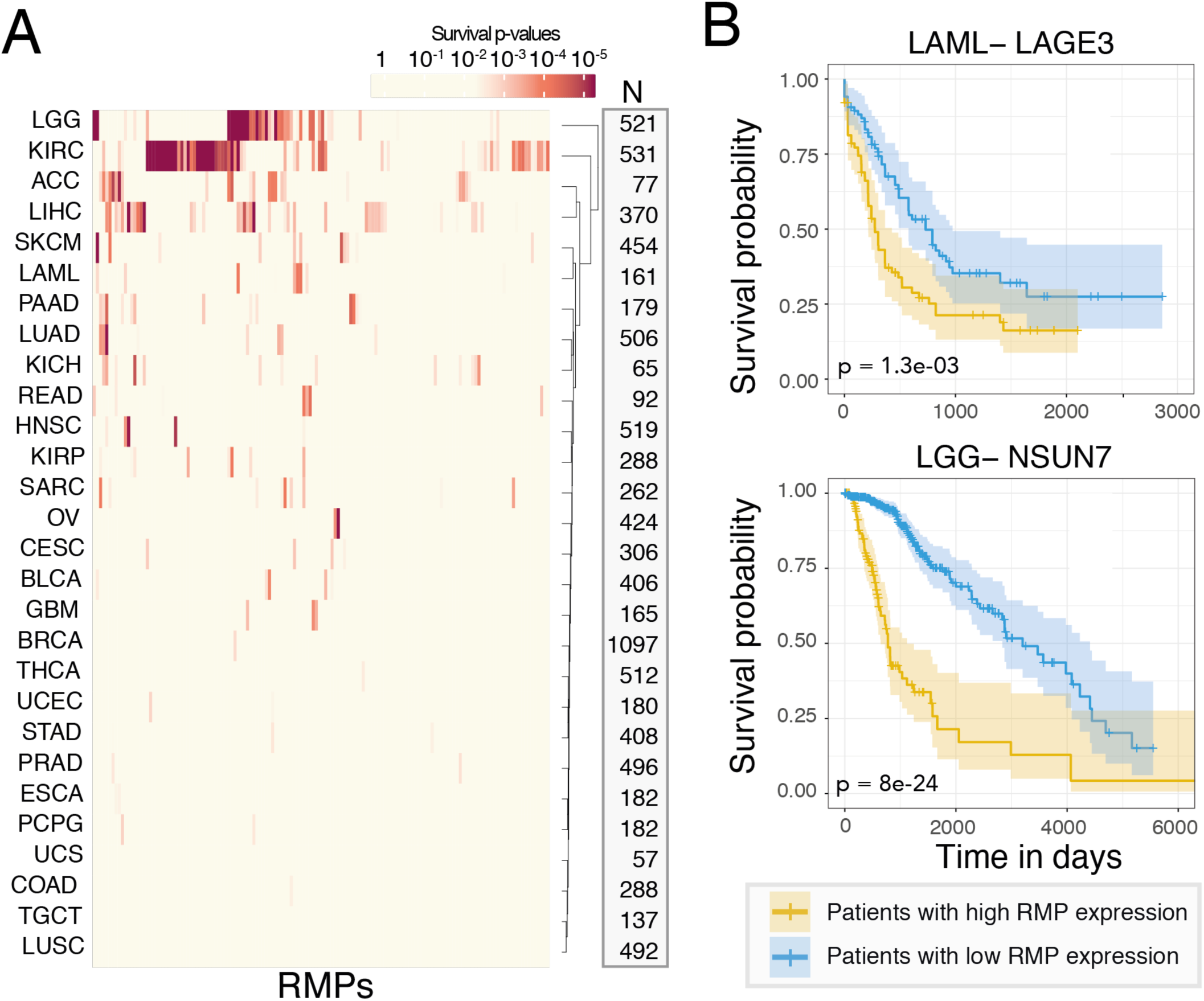
Prognostic value of RMPs expression levels in different cancer types. **(A)** Heatmap of survival p-values of 146 RMPs across 28 cancer types. Survival p-values are calculated by comparing the prognosis of patients that express high (upper 50%) versus low (lower 50%) RMP levels. “N” column shows the number of patients included for the analysis of each cancer type. **(B)** Individual examples of survival plots where the expression levels of the RMP are predictive of cancer prognosis. P-values have been calculated by comparing the survival between patients expressing high levels (yellow, top 50%) versus low expression levels (blue, bottom 50%).

Surprisingly, we found that FTO expression levels are not significantly correlated with patient survival in LAML, despite studies are currently focusing on this cancer type to test FTO inhibitors (**Table S7**). By contrast, LAGE3 expression levels were significantly correlated with patient survival in LAML (**Figure 5B**). Among all the RMP-cancer pairs studied, we identified NSUN7 as the top-ranked RMP in terms of prediction of lower grade glioma (LGG) patient survival (*p*=8e^-24^); although its biological role still remains uncharacterized. Future research will be needed to functionally dissect the role that NSUN7 plays in glioma, as well as to decipher why its expression levels are highly predictive of patient survival.

## DISCUSSION

Over the past decade, systematic efforts to detect and map RNA modifications have boosted the new field of epitranscriptomic research. Many proteins are involved in the writing, reading and erasing of RNA modifications but their roles in tumorigenesis and potential as therapeutic targets remain largely uncharacterized. To bridge this gap, here have compiled a list of 146 human RNA modification-related proteins (RMPs) (**Table S1**), and have analyzed the evolutionary history and gene expression patterns of 90 RMPs across 32 mammalian tissues, 10 species, 5 cell types and 13,358 tumor-normal paired cancer samples.

Through this analysis, we identify a large amount of duplication events in multiple RNA modification families (**Figure 1**), and find that duplications are often accompanied by the acquisition of restricted tissue expression patterns and/or change in its RNA target specificity. We find that the majority of tissue-restricted RMPs are in fact testis-enriched (**Figure 2**), suggesting that certain RMPs might play a pivotal role in sperm formation and maturation. Indeed, deletion of testis-enriched genes such as NSUN7, ADAD1 or HENMT1 leads to male sterility [5,55,56,60]

At the beginning of spermatid elongation, nuclear condensation starts, and consequently the transcriptional machinery is shut down. Therefore, to provide proteins for the following maturation steps of sperm assembly, mRNAs have to be premade in spermatocytes and round spermatids, before nuclear condensation happens, and translationally repressed until needed [49–52]. Chemical RNA modifications provide an ideal platform to achieve the fine regulation that is required upon transcriptional shutdown, determining which RNAs are expressed, repressed, or undergo decay [76]. In this regard, previous work has shown that METTL3/METTL14 mediated m^6^A modification is dynamically regulated in spermatogenesis [34]. Similarly, piRNA molecules in germ line cells are tightly regulated by HENMT1, via 2’-O-methylation of their 3’ends [5]. Here we show that a vast proportion of RMPs are dynamically regulated during spermatogenesis as well as during sperm maturation in the epididymis, and as such, may be involved in the regulation and decay of specific transcripts that occur during sperm formation and maturation (**Figure 3**).

Recent works shown that specific RNA modifications are essential for the transmission of paternal diet-induced phenotypes intergenerationally [53]. Here we identify two RMPs (TRDMT1 and METTL1) whose expression is significantly enriched in epididymis (**Figure S3**), one of which (TRDMT1) was recently shown to be involvedin the transmission of diet-induced paternal phenotypes across generations[53]. Whether METTL1 plays a role in intergenerational inheritance is yet to be deciphered; however, recent insights showing its role in miRNA maturation [58] suggest that this enzyme might be playing a role in miRNA-acquired inheritance of information.

In the last few years, several studies have placed RNA modifications in the forefront of cancer research[36,38,64,66,77], mostly focused on the machinery responsible for writing and erasing m6A modifications. For many years, FTO was thought to be of special interest due to its association with obesity [78]. However, later studies proved this genome-wide association to be false [79], and that the single nucleotide variant present in the FTO intron was in fact associated with the activity of neighbouring genes [79].

Nonetheless FTO kept receiving special attention due to its perceived activity as an eraser of N6-methyladenosine (m6A) [27], the most frequent type of RNA modifications present in mRNAs. However, this is now thought to be incorrect, as later studies showed that FTO is in fact an eraser of N6,2’O-methyladenosine (m6Am), which is much less abundant in mRNAs [28,80]. Similarly, FTO has been proposed to constitute a promising target for antitumor therapies [38,81,82]. While FTO has been shown to play an important role in leukaemia [81], it is possible that additional RMPs such as HENMT1, which is drastically dysregulated in this cancer type, might constitute a better drug target to inhibit leukemogenesis (**Figure 4**).

Here we show that the expression of 40 RMPs is significantly altered in tumor samples, relative to their matched normal samples (**Table 1** and **Figure 4**). Moreover, we identify two enzymes, LAGE3 and HENMT1, as the top recurrently up-regulated RMPs across cancer types. Surprisingly, these proteins have so far received little attention in cancer research studies. LAGE3 mutations are known to cause multiple human diseases, including nephrotic syndrome and microcephaly [83]; however, its role in tumorigenesis and cancer progression is yet to be determined. Future work will be needed to decipher the biological role of LAGE3 and HENMT1 in cancer, as well as its potential use as a target for diagnostic and prognostic purposes.

## CONCLUSIONS

Our analyses reveal an unanticipated heterogeneity in the expression patterns of RMPs across healthy mammalian tissues, with an over-representation of testis-specific RMPs, many of which are essential for sperm formation and maturation, and in some cases, are required for the transmission of epigenetic information across generations. In addition, we uncover a large proportion of dysregulated RMPs in multiple cancer types, and show that several RMPs are dysregulated to a much larger extent than commonly studied m6A modification pathway, stressing the need to extend the epitranscriptomic drug targeting strategies to additional RNA modification enzymes. Now that novel transcriptome-wide tools to map additional RNA modifications have been recently made available, the community can repurpose antitumoral strategies to those RNA modification pathways that are in fact most significantly dysregulated in each cancer type.

## METHODS

### Compilation of human RNA modification-related proteins (RMPs)

An initial list of human methyltransferases, deaminases and pseudouridylases was obtained by merging the lists available in the MODOMICS database (http://modomics.genesilico.pl/) and from a recently published review [13]. These lists were further initially completed with candidate genes by the addition of annotated proteins on Uniprot [42]. For each of these proteins, hidden Markov model (HMM) profiles of the corresponding PFAM catalytic domains were retrieved (**Table S8**) by querying the PFAM database (https://pfam.xfam.org/). Each HMM profile then was used to query human proteome using hmmsearch function from HMMER software v.3.2.1 (http://hmmer.org/). Proteins above default threshold we kept as candidate RMW proteins (**Table S9**). Related information for each of these proteins (modification type, target RNA, localization) was extracted from Uniprot, as well as from relevant literature [42]. Additional tRNA writer proteins were gathered from a recent study matching tRNA modifications to their writers [84]. Readers, erasers and non-catalytic subunit proteins were obtained from annotated Uniprot genes as well as from published literature [85]. APOBEC3G and APOBEC3A were included in the analyses due to recent literature showing their deamination activity on RNA molecules *in vivo* in addition to acting on DNA [86,87]

### Phylogenetic analysis

We first built a set of representative eukaryotic species, by choosing one species for each major phylogenetic clade for which complete proteomes were available. Our final list of representative species consisted of 25 complete proteomes from UniProt [42], which included 23 eukaryal species, as well as 2 outgroups (1 bacteria and 1 archaea) **(Table S10**). For each proteome and RMW, we performed HMM-based searches, as described above. Candidate orthologs were manually curated to ensure that we did not miss any ortholog in our analysis, which resulted in a final table of RMW ortholog proteins **(Table S11)**. For each curated ortholog dataset, multiple sequence alignments were built using MAFFT with G-INS-1 method [88]. Alignment files were used to construct maximum-likelihood phylogenetic tree using IQ-Tree with bootstrapping (n=5000) [89]. Consensus trees were visualized using FigTree v 1.4.4 [90] and used to identify the duplication events (**Table S2**).

### Tissue specificity analysis

Human mRNA expression levels (TPM-Transcripts Per Kilobase Million) for each of the 146 human RMPs were downloaded from the Genotype Tissue Expression (GTEx) dataset [45], version v7, as well as from the Human Protein Atlas (HPA)[91]. Three GTEX tissues (whole blood, transformed lymphocytes and transformed fibroblasts) were discarded from downstream analyses, as these have been previously considered as outliers that can bias the analyses [45] or are not normal tissues of the human body. mRNA expression levels for adult mouse tissues (TPM, Transcripts Per Kilobase Million) were obtained from ENCODE3 (http://genome.crg.es/~avlasova/ENCODE3/STAR-RSEM/). For each dataset (HPA, GTEx, ENCODE), we log transformed the TPM values after the addition of a pseudocount. To determine which genes were tissue-specific, we compared the expression levels of RMP in a given tissue to the median expression levels of RMPs across all tissues. We then calculated residuals (using *rlm* function), which we refer to as “tissue-specificity score” (TS), for each RMP to the regression line of each tissue. An RMP was considered tissue-specific if their TS was greater than 2.5 standard deviation (SD), as previously described [48], which, in a normal distribution of the standardized residuals, equals to the region outside of the 97.9 percentiles.

### RMP expression analysis across tissues in amniote species

mRNA expression levels of 12 amniote species (human, chimpanzee, bonobo, gorilla, orangutan, rhesus macaque, mouse, gray-short tailed opossum, platypus and chicken) were obtained from GSE30352 [92]. Normalized RPKM values of constitutive exons for both amniote and primate orthologs were used for downstream analyses. Heatmaps of the log transformed (with a pseudocount) and row (gene) z-scaled tissue-wide mRNA expression values were built using *complex heatmap* R package. PCA analysis was performed using *prcomp* function of R and plots of scores (amniote and primate tissues) and loadings (orthologue genes) were plotted for the first two principal components using *ggplot* R package.

### Analysis of RMPs expression during spermatogenesis

Processed spermatogenesis data [54] was extracted from GSE112393. Input data was used to perform k-means clustering of RMPs based on their expression profiles in different sperm cell populations. The optimal number of clusters was calculated by plotting the within groups sum of squares by number of clusters extracted using k-means function in R, following criteria used by Scree’s test. Heatmaps were built using *complex heatmap* R package. Violin plots were built using the *ggplot* R package.

### Immunohistochemistry

Testis and epididymis from C57BL/6J mice were fixed overnight at 4°C with neutral buffered formalin (HT501128-4L, Sigma-Aldrich) and embedded in paraffin. Paraffin-embedded tissue sections (3 µm in thickness) were air dried and further dried at 60°C overnight. Immunohistochemistry was performed using The Discovery XT Ventana Platform (Roche). Antigen retrieval was performed with Discovery CC1 buffer (950-500, Roche). Primary antibodies rabbit polyclonal anti-NSUN2 (20854-1-AP, Proteintech), rabbit polyclonal anti-NSUN7 (PA5-54257, Thermo Fisher Scientific), rabbit polyclonal anti-HENMT1 (PA5-55866, Thermo Fisher Scientific), and rabbit polyclonal anti-METTL14 (HPA038002, HPA038002) were diluted 1:1000, 1:100, 1:150 and 1:2000 respectively with EnVision FLEX Antibody Diluent (K800621, Dako, Agilent) and incubated for 60 min. Secondary antibody OmniMap anti-rabbit HRP (760-4311) was incubated for 20 min. Detection of the labelling was performed using the ChromoMAP DAB (760-159, Roche). Sections were counterstained with hematoxylin (760-2021, Roche) and mounted with Dako Toluene-Free Mounting Medium (CS705, Agilent) using a Dako CoverStainer (Agilent). Specificity of staining was confirmed with a rabbit IgG, polyclonal Isotype Control (ab27478, Abcam). Brightfield images were acquired with a NanoZoomer-2.0 HT C9600 digital scanner (Hamamatsu) equipped with a 20X objective. All images were visualized with a gamma correction set at 1.8 in the image control panel of the NDP.view 2 U123888-01 software (Hamamatsu, Photonics, France). Mice samples were collected, prepared as paraffin blocks, sliced and stained at the IRB Histopathology Facility. Negative controls for each antibody were also included, which showed no staining (**Figure S6**).

### Analysis of RMP expression in tumor-normal paired human datasets

TPM expression values were downloaded from the UCSC XENA Project, which contains the TCGA and GTEX RNA Seq data that is processed together to provide more reliable expression analysis with tumor and normal samples [69]. We discarded CHOL, THYM and DLBC tumor-normal tissue pairs due to lack of proper control of normal tissue (low number of patients) in these cancer types. Data was transformed into log2(TPM+1) for downstream analyses. For the log2(FC) analyses, we calculated the difference between median log2 expression levels between tumor and normal datasets, for each cancer type and RMP. For dysregulation analysis, we calculated the residuals (using *rlm* function in R) for all of the gene expression in a given tumor tissue and normal tissue pair, which has been previously termed as ‘dysregulation score’ (DS) [48]. We set the threshold of significance DS at 2.5 standard deviations (SD) as previously described [48], which, in a normal distribution of standardized residuals, equals to the region outside of the 97.9 percentiles. We then extracted the dysregulation scores of the RMPs and used it for further downstream analyses. For heatmap representations, dysregulation scores were scaled and centered, and the final heatmap was built using *complex heatmap* R package. Scatter plots of median log2 expression values for all genes in tumor-normal paired data were built using the *ggplot* R package, highlighting RMPs in black, significantly dysregulated RMPs in red (up-regulated in tumor) and blue (down-regulated in tumor), and non-RMP proteins were depicted in grey.

### Survival Analyses

Survival phenotypes were downloaded from the XENA Platform, using the “TCGA TARGET GTEX” cohort [69]. In order to analyse the survival data, we first determined patients that have “high” (upper 50% relative to average expression) and “low” (lower 50% relative to average expression) expression of a specific gene, and matched these patients with their overall survival information. We then used the *survminer* R package to plot survival curves for each gene and every cancer type, as well as to extract the survival p-values. P-values were transformed by inversion and subsequent log-transformation with a pseudocount [log(1/p+1)]. Heatmap of the survival p-values was built using *complex heatmap* R package. Transformed survival p-values were visualized using *ggplot*.

## DECLARATIONS

### Code availability

All scripts used in this work will be made publicly available, and will be found at https://github.com/obegik/RNAModMachinery, once the paper is published.

### Data availability

All datasets used to build the figures, as well as intermediate analysis files (alignment files, maximum likelihood trees, scatter plots of tissue-specificity, barplots of amniote and primate ortholog expressions, scatter plots of tumor vs normal tissues, boxplots of individual expression of RMPs in tumor-normal paired tissues and survival plots) and raw IHC scans will be made publicly available when the paper is published.

### Authors’ contributions

OB performed all the analyses. MCL contributed to IHC experiments as well as to the corresponding figures. HL and JMR contributed to code and suggestions for the data analysis. OB made all figures, with the contribution from MCL. OB and EMN interpreted the results. EMN conceived the project. EMN and JSM supervised the work. OB and EMN wrote the manuscript, with input from all authors.

## Acknowledgements

We thank all the members of the Mattick and Novoa labs for their valuable insights and discussion. The results shown here are in whole or part based upon data generated by the TCGA Research Network.

